# Deciding between one-step and two-step irreversible inhibition mechanisms on the basis of “*k*_obs_” data: A statistical approach

**DOI:** 10.1101/2020.06.08.140160

**Authors:** Petr Kuzmič

## Abstract

This paper describes an objective statistical approach that can be used to decide between two alternate kinetic mechanisms of covalent enzyme inhibition from kinetic experiments based on the standard “*k*_obs_” method. The two alternatives are either a two-step kinetic mechanism, which involves a reversibly formed noncovalent intermediate, or a one-step kinetic mechanism, proceeding in a single bimolecular step. Recently published experimental data [Hopper *et al.* (2020) *J. Pharm. Exp. Therap.* **372**, 331–338] on the irreversible inhibition of Bruton tyrosine kinase (BTK) and tyrosine kinase expressed in hepatocellular carcinoma (TEC) by ibrutinib (PCI-32765) and acalabrutinib are used as an illustrative example. The results show that the kinetic mechanism of inhibition was misdiagnosed in the original publication for at least one of the four enzyme/inhibitor combinations. In particular, based on the available *k*_obs_ data, ibrutinib behaves effectively as a one-step inhibitor of the TEC enzyme, which means that it is not possible to reliably determine either the inhibition constant *K*_i_ or the inactivation rate constant *k*_inact_, but only the covalent efficiency constant *k*_eff_ = *k*_inact_/*K*_i_. Thus, the published values of *K*_i_ and *k*_inact_ for this system are not statistically valid.

## 1. Introduction

Covalent enzyme inhibitors form a chemical bond between a reactive functional group on the molecule and the target protein. Even though there are examples of reversible covalent inhibitors [1], the formation of the covalent bond is usually irreversible. This property has been used to design therapeutic agents purposely inhibiting overactive enzymes. Probably the most well known covalent inhibitor drug in this category is Aspirin, an irreversible inhibitor of cyclooxygenase (COX).

From the physical point of view, the irreversible inhibition mechanism must include two separate steps. In the first step, the enzyme and inhibitor associate reversibly to form a noncovalent complex. In the second step, the noncovalent complex undergoes an irreversible transformation. This two-step molecular mechanism can be schematically represented as E + I ⇌ E · I → EI. However, from the point of view of formal kinetic analysis, many enzyme inhibitors outwardly behave as if the initial noncovalent complex were absent. In those cases the experimental data give the impression that the two-step mechanism is somehow fused into a single irreversible step, E+ I → EI. The occurrence of the one-step irreversible inhibition mechanism has been described as “nonspecific affinity labeling” [2, sec. 7.2.1], involving generic inhibitors similar to iodoacetate and N-ethyl maleimide. These inhibitors are assumed to have negligibly low initial binding affinity and to indiscriminately modify “many amino acid residues on the enzyme molecule” [3, sec 9.1]. The lack of selectivity presumably explains the one-step kinetic behavior. In contrast, highly selective inhibitors that precisely target the enzyme’s active site are assumed to follow the two-step kinetic mechanism [2, 3]. However, anecdotal evidence from this investigator’s workshop suggests that numerous highly specific irreversible inhibitors also show one-step kinetics, specifically in kinase inhibition assays.

The full understanding of this difficult subject has been hampered by the fact that the available literature on irreversible enzyme inhibition does not offer sufficient advice on how to properly distinguish between candidate kinetic mechanisms. The discussion usually starts from the premise that that kinetic mechanism (either one-step or two-step) is known in advance. Mathematical formulas are then presented (for example, a certain linear equation or, alternately, a certain hyperbolic equation) that can be used to analyze experimental data conforming to each inhibition mechanism. What the textbook literature does not explain is how we can tell, on the basis of a given set of experimental data, which kinetic mechanism is most likely to be operating. This question is the main subject of the present report. The data sets being investigated here are of a certain type that is frequently encountered in drug discovery research. Raw experimental data on the inhibition of BTK and TEC kinase enzymes by ibrutinib and acalabrutinib, recently published by Hopper *at al.* [4], serve as an illustrative example.

## 2. Methods

This section describes the theoretical, mathematical, and statistical methods that were used to analyze the experimental data originally published in ref. [4]. All computations were performed by using the software package DynaFit [5, 6]. Explanation of all mathematical symbols is given in the Appendix, see *Table A.1* and *Table A.2*.

### 2.1. Kinetic mechanisms of irreversible inhibition

In this report we will consider in various contexts the kinetics mechanisms of substrate catalysis and irreversible inhibition depicted in *Figure 1*. For details see ref. [7], which also contains an in-depth discussion of the important difference between the equilibrium dissociation constant *K*_i_ and the inhibition constant *K*_I_.

**Figure 1:**
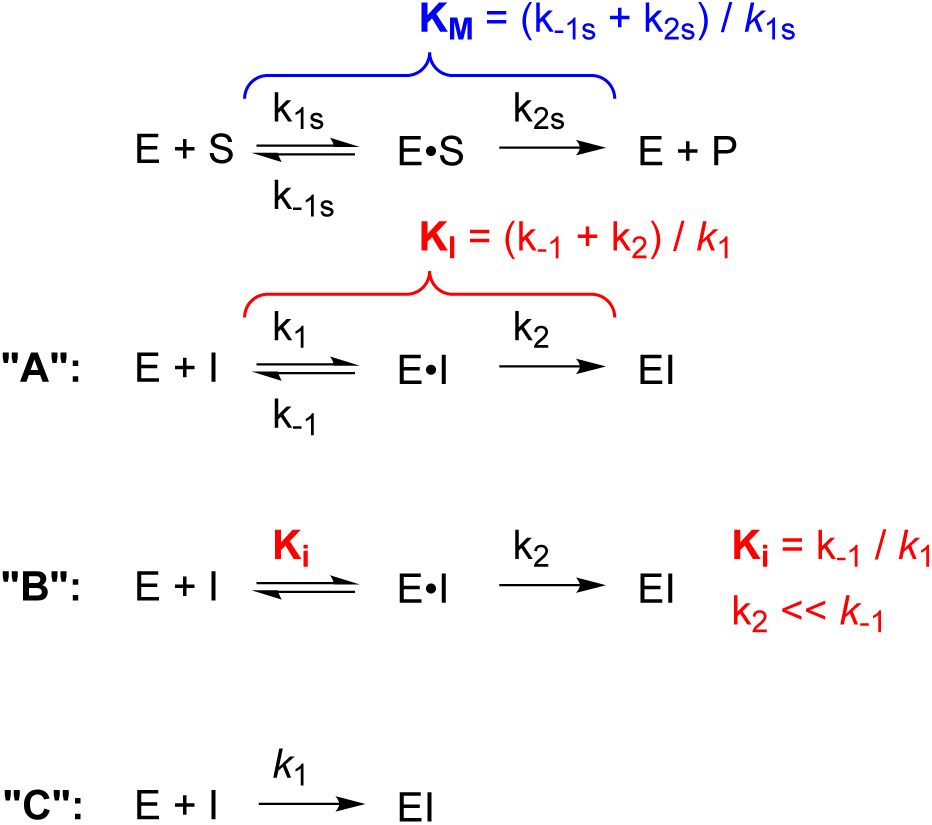
Kinetic mechanisms of substrate catalysis (top) and covalent inhibition (mechanisms **A – C**).

### 2.2. Mathematical models

Under a number of simplifying assumptions first introduced into covalent inhibition analysis by Kitz & Wilson [8], Tian & Tsou [9] showed that the time course of a irreversible inhibition assay can be described by a rising exponential curve according to Eqn (1), where *F* is some experimental signal such as fluorescence intensity; *F*_0_ is the experimental signal observed at time zero (i.e., a baseline signal as a property of the instrument); *V*_*i*_ is the initial reaction rate in arbitrary instrument units, observed at the given inhibitor concentration [I]_0_; and *k*_obs_ is an apparent first-order rate constant.

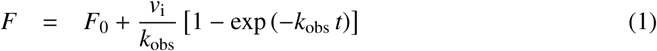

Assuming that the two-step kinetic mechanism **B** is operating, the apparent first-order rate constant *k*_obs_ depends on the inhibitor concentration according to the hyperbolic Eqn (2), where 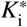 is the apparent inhibition constant and *k*_inact_ is the first-order inactivation rate constant. In the case of the one-step mechanism **C**, *k*_obs_ depends on [I]_0_ according to the linear equation Eqn (3), where the slope parameter 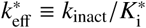 is the apparent covalent efficiency constant.

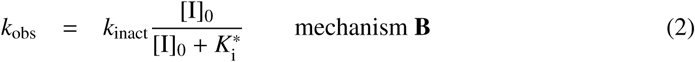

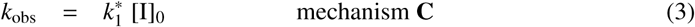

Assuming that the inhibitor is kinetically competitive with the substrate, the experimentally observable apparent values of 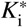 (two-step mechanism **B**) or 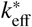 (one-step mechanism **C**) are related to their true values *K*_i_ and *k*_eff_ as shown in Eqns (4)–(5), where *K*_M_ is the Michaelis constant.

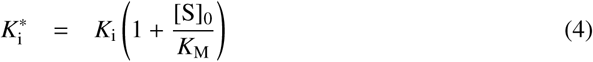

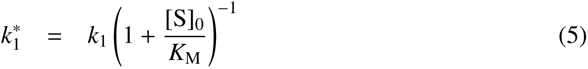

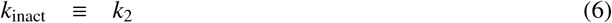

### 2.3. Model selection methods

We propose a multi-pronged approach relying on four independent statistical methods, resulting in four independent model acceptance criteria. The more complex (two-step) kinetic mechanism **B** will be accepted in favor of the simpler (one-step) mechanism **C** only if all four types of statistical model selection criteria are satisfied at the given confidence level (for example, 99%).

#### 2.3.1. Profile-t method for confidence intervals

For any given *k*_obs_ vs. [I]_0_ data set, we will tentatively accept the two-step mechanism **B** in favor of the one-step mechanism **C** only if the upper limits for both the inhibition constant 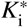 and the inactivation rate constant *k*_inact_ are well defined by the available data. These upper limits (if they do exist) can be found at the given confidence level, e.g. 99%, by using the profile-*t* method [10, 11].

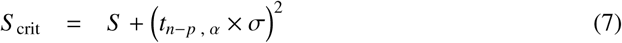

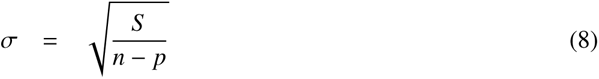

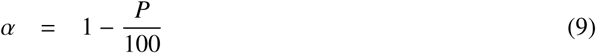

Briefly, the least-squares regression analysis is repeated at multiple *search points* where either 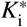 or *k*_inact_ is held constant while the remaining parameter is optimized. The systematic search for the confidence interval limits proceeds until the residual sum of squares increases from its best-fit value, *S* in Eqn (7), to the *critical value*, *S* _crit_. The critical value *S* _crit_ is larger than the best-fit value *S* by a specific increment defined by the additive factor in Eqn (7), where *t*_*n−p*, *α*_ is the Student-*t* statistic with *n − p* degrees of freedom at the confidence level *α*; *n* is the number of data points and *p* is the number of adjustable model parameters. A suitable search algorithm, implemented by the software package DynaFit [6], is described in ref. [10, p. 302], Appendix A3.5.1.

#### 2.3.2. Akaike Information Criterion and Akaike weights

The hyperbolic fitting Eqn (2) contains two adjustable model parameters (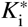 and *k*_inact_) whereas the linear Eqn (3) contains only one parameter 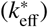. Therefore, it is not possible to decide which model is “better” solely on the basis of the residual sum of squared deviations. A suitable correction for the difference in the number of parameters is provided by information-theoretic model selection criteria, such as the Akaike Information Criterion (AIC) defined by Eqn (10); see refs. [12–14] for details.

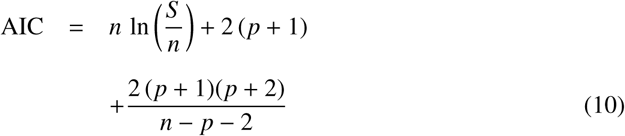

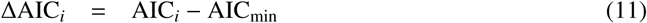

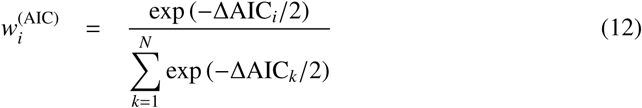

In Eqn (10), *S* is the best-fit residual sum of squares; *n* is the number of data points; and *p* is the number of adjustable parameters. The differential Akaike Information Criterion for the *i*th fitting model, ∆AIC_*i*_, is computed according to Eqn (11), where AIC_min_ is the lowest AIC value. Finally the *Akaike weight* for the *i*th model is defined by Eqn (12) [12, p. 75]. In this work we will accept the more complex two-step inhibition mechanism if the Akaike weight associated with Eqn (2) is larger than 0.99 or 0.95, corresponding to 99% or 95% likelihood.

#### 2.3.3. Bayesian Information Criterion and Bayesian weights

The Bayesian information criterion, BIC, defined by Eqn (13), is a complementary information-theoretic alternative to the AIC [13, 14]. The *Bayesian weight* is defined by Eqn (15).

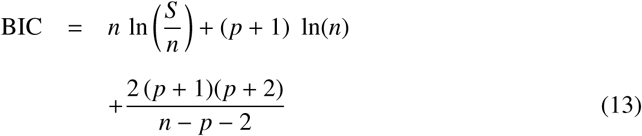

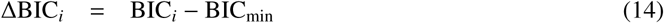

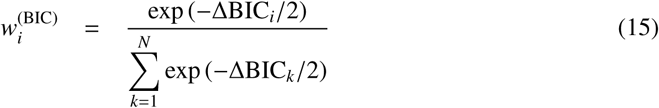

The Bayesian weight for any given fitting model is a number between zero and one and measures the statistical likelihood that the given candidate is the true model. We will accept the two-step inhibition mechanism **B** represented by Eqn (2) if the corresponding Bayesian weight is larger than 0.95 or alternately 0.99.

#### 2.3.4. F-test for nested models

Mannervik [15] described a model selection procedure for two candidate fitting equations in the special case where the equations represent two *nested models*. Note that Eqn (2) of the general form *Y* = *aX*/(*b* + *X*) is an algebraic extension of Eqn (3), of the general form *Y* = *aX*. In that sense Eqns (2)–(3) represent nested models.

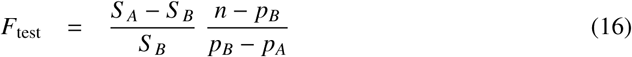

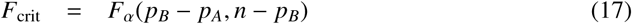

The general formula used for model selection is shown in Eqn (16), where subscript *A* represents the reduced (or smaller) fitting model and subscript *B* represents the full (or lager) fitting model. In Eqn (16), *n* is the number of experimental data points; *S*_*A*_ and *S*_*B*_ are the residual sums of squares; and *p*_*A*_ and *p*_*B*_ represent the number of adjustable model parameters. The ratio *F*_test_, computed according Eqn (16), is compared with the upper critical value *F*_crit_ of the *F* distribution for *p*_*B*_ − *p*_*A*_ and *n* − *p*_*B*_ degrees of freedom at the probability level *α*. If the variance ratio *F*_test_ is larger than the critical value *F*_crit_, the fitting equation with the larger number parameters is accepted as plausible.

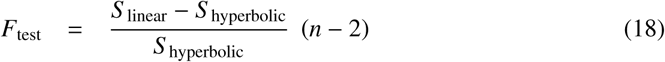

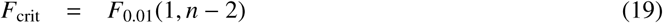

In the specific case of deciding between the linear Eqn (3) and the hyperbolic Eqn (2), the *F*_test_ ratio is computed according to Eqn (18), whereas the upper critical value *F*_crit_ at 99% probability level (*α* = 0.01) is computed according to Eqn (19).

## 3. Results

In the remainder of this report, the *k*_obs_ vs. [I]_0_ data sets originally published in ref. [4] are numbered according to the numbering scheme displayed in *Table 1*.

**Table 1:**
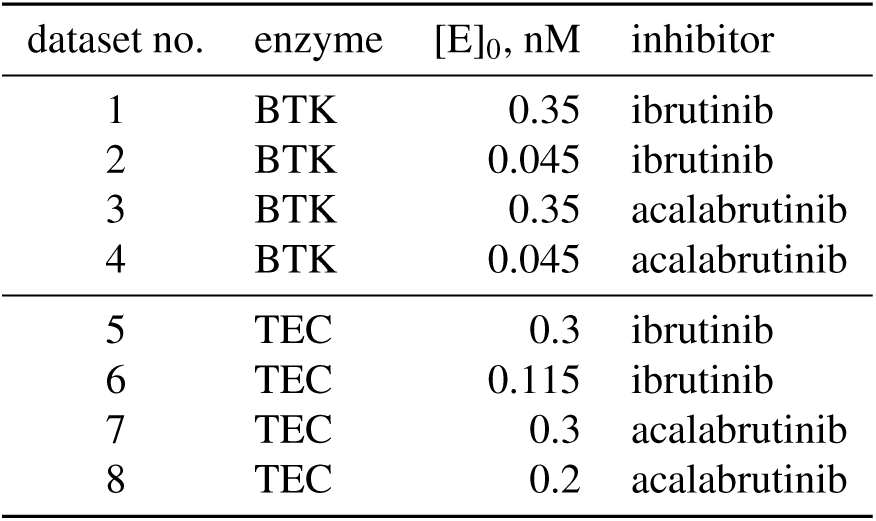
Data set numbering corresponding to Supplemental Tables and Supplemental Figures in ref. [4].

### 3.1. Profile-t confidence intervals

The results of statistical model discrimination based on the profile-*t* method are summarized in *Table 2* for the apparent inhibition constant 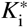 and in *Table 3* for the inactivation rate constant *k*_inact_. The dash (–) indicates that that upper limit of the given parameter could not be determined at the given confidence level.

**Table 2:**
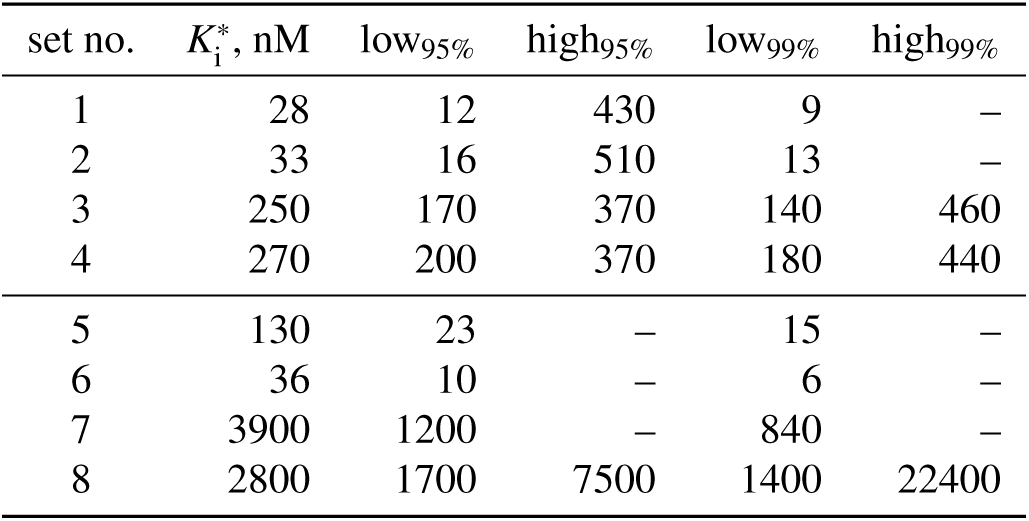
Results of model discrimination analysis based on the profile-*t* method: non-symmetrical confidence intervals for the apparent inhibition constant 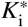. For details see text.

| set no. | $K_i^*$, nM | low<sub>95%</sub> | high<sub>95%</sub> | low<sub>99%</sub> | high<sub>99%</sub> |
| --- | --- | --- | --- | --- | --- |
| 1 | 28 | 12 | 430 | 9 | – |
| 2 | 33 | 16 | 510 | 13 | – |
| 3 | 250 | 170 | 370 | 140 | 460 |
| 4 | 270 | 200 | 370 | 180 | 440 |
| 5 | 130 | 23 | – | 15 | – |
| 6 | 36 | 10 | – | 6 | – |
| 7 | 3900 | 1200 | – | 840 | – |
| 8 | 2800 | 1700 | 7500 | 1400 | 22400 |

**Table 3:**
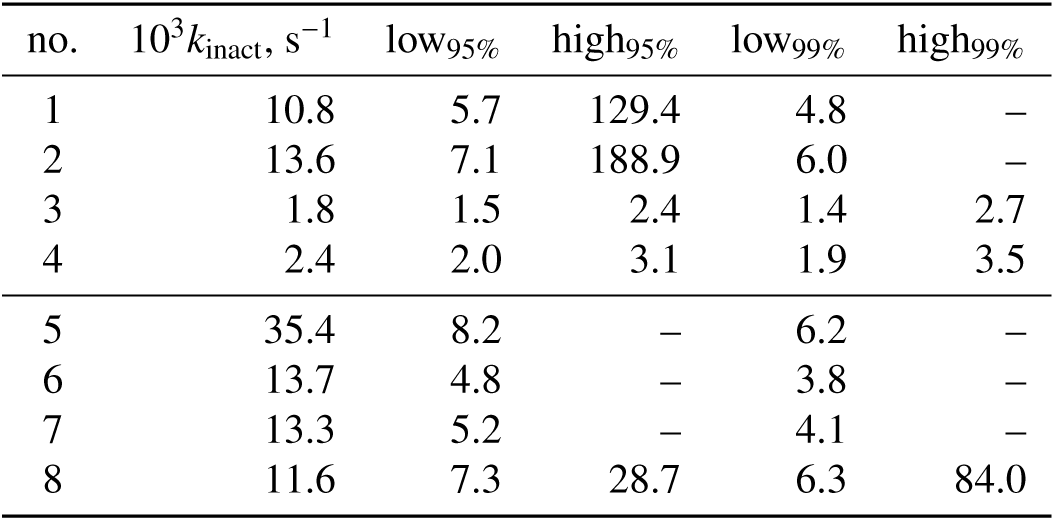
Results of model discrimination analysis based on the profile-*t* method: non-symmetrical confidence intervals for the inactivation rate constant *k*_inact_. For details see text.

The upper limit results listed in *Table 2* and *Table 3* can be interpreted mechanistically as follows. At any given confidence level (95% or 99%), we can scan down the column labeled “high” and check to see whether the upper limit of 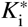 or *k*_inact_ is defined. If the “high” column does contain a defined upper limit, we can conclude that the hyperbolic Eqn (2) corresponding to two-step mechanism **B** provides a good description of that data set. In contrast, if the “high” column does not contain a defined value for the upper limit, we must conclude that the linear Eqn (3) corresponding to the one-step mechanism **C** provides a better description of the available data. The results show that only the inhibition of BTK by acalabrutinib conclusively proceeds by the two-step mechanism, because the upper limits of confidence intervals for 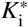 and *k*_inact_ are defined for both available data sets (experiments no. 3 and 4) and also at both confidence levels (95% and 99%).

The only other instance of a defined upper limit for either 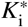 or *k*_inact_, at both confidence levels, was observed in data set no. 8 (acalabrutinib inhibition of TEC, second replicate). However, at the 99% confidence level the high/low ratio for 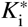 is quite high, 22400/1400 = 16, spanning more than an order of magnitude. Under such circumstances we might suspect that the defined upper limit of the confidence interval might have arisen by random chance, rather than being firmly supported by the data. The results for ibrutinib inhibition of BTK (data set no. 1 and 3) are also ambiguous, in the sense that the upper limits of the confidence intervals are defined at the 95% confidence level but undefined at the 99% confidence level. Note again that at the 95% confidence level the confidence intervals for ibrutinib inhibition of BTK are exceedingly wide. For example, for data set no. 1, the high/low ratio was 430/12 = 36. The two examples below provide graphical illustrations of two distinct types of kinetic behavior, according to the results listed in *Table 2* and *Table 3*.

#### Example 1: Dataset No. 4

In this example representing acalabrutinib inhibition of BTK, the upper limits of the confidence intervals for 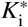 and *k*_inact_ are clearly defined the 99% confidence level, signifying a strong support for the two-step inhibition mechanism **B**. The results are shown in *Figure 2*.

**Figure 2:**
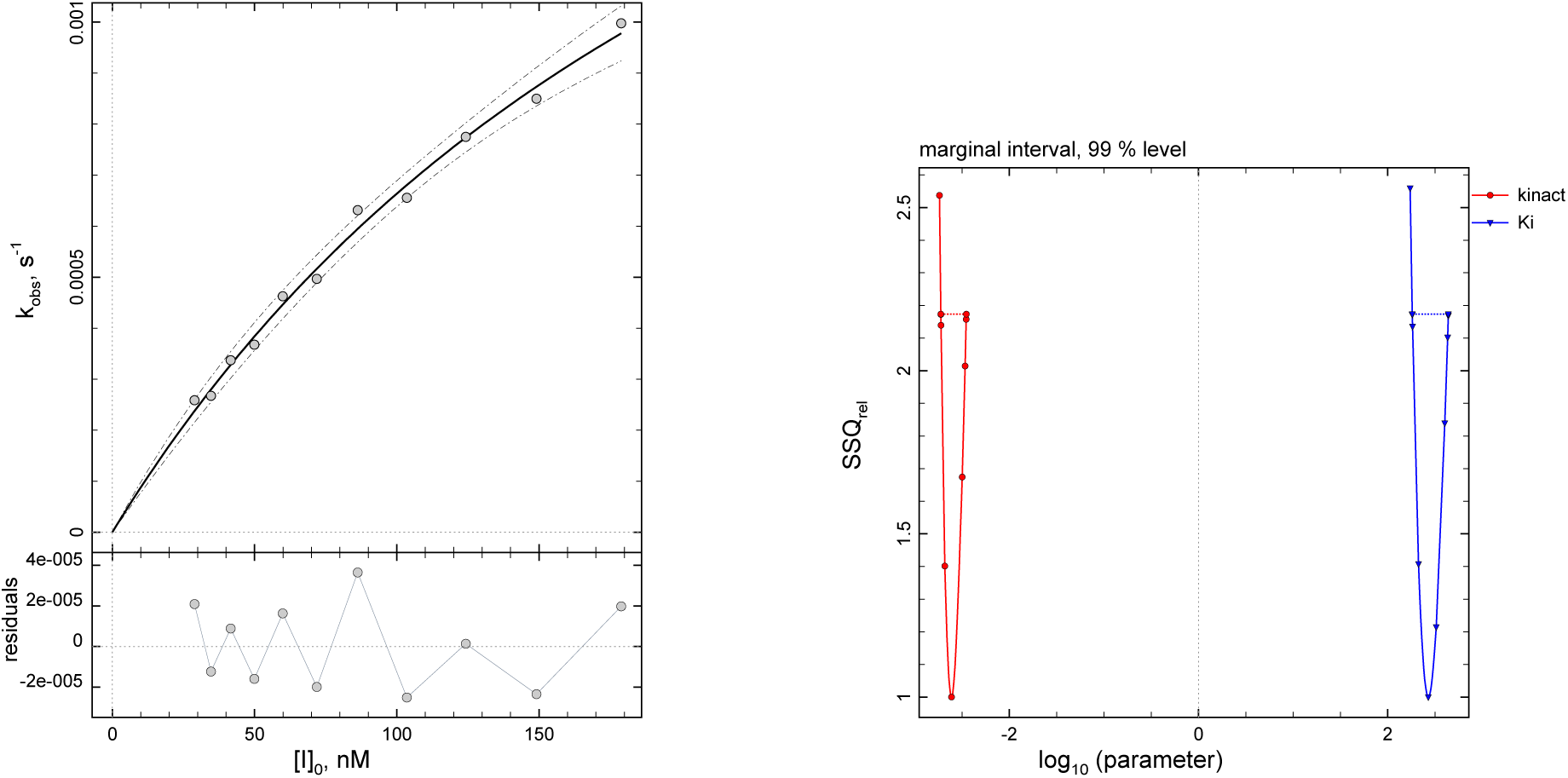
Results of fit to the two-step hyperbolic Eqn (2), data set no. 4. **Left:** Experimental data (circles), best-fit model curve, and the residual plot. **Right:** Confidence interval profiles for the kinetic constants 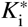 and *k*_inact_.

In *Figure 2*, the top panel displays the overlay of the experimental data (circles) on the best-fit model curve and the corresponding residual plot. The dashed envelope curves in the upper panel represent the model confidence bands. The bottom panel shows the results of the confidence interval search for either the inhibition constant 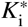 (blue profile curve on the right) or the inactivation rate constant *k*_inact_ (red profile curve on the left). The horizontal axis in the bottom panel of *Figure 2* represents the logarithm of the search-point values (blue triangles for 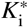 and red circles for *k*_inact_) that were visited according to the profile-*t* search algorithm [10]. The vertical axis represents the relative sum of squares, that is, a ratio of the best-fit residual squares corresponding to the given search point, divided by the best-fit residual squares corresponding to the optimal point. The thin dotted horizontal lines correspond to the critical value of the residual squares defined by Eqn (7). Both confidence interval profiles depicted in *Figure 2* have an approximately parabolic shape, are relatively narrow (see *Table 2* and *Table 3*), and, most importantly, have a well defined upper limit.

#### Example 2: Dataset No. 6

In this example representing ibrutinib inhibition of TEC, the upper limits of confidence intervals for 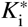 and *k*_inact_ are undefined at the given confidence level, signifying that the two-step hyperbolic equation Eqn (2) is *not* supported by the data and therefore we must chose the simpler linear fitting Eqn (2) for this data set. The results are shown in *Figure 3*. The bottom panel *Figure 3* shows the results of the confidence interval search for either the inhibition constant 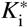 or the inactivation rate constant *k*_inact_. In contrast with the Example 1, the confidence interval profiles are non-parabolic and show only a very shallow minimum. Most importantly, the upper limits of the confidence intervals for either 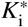 or *k*_inact_ are undefined, which signifies that the two-step hyperbolic equation Eqn (2) is not an adequate theoretical model for data set no. 6.

**Figure 3:**
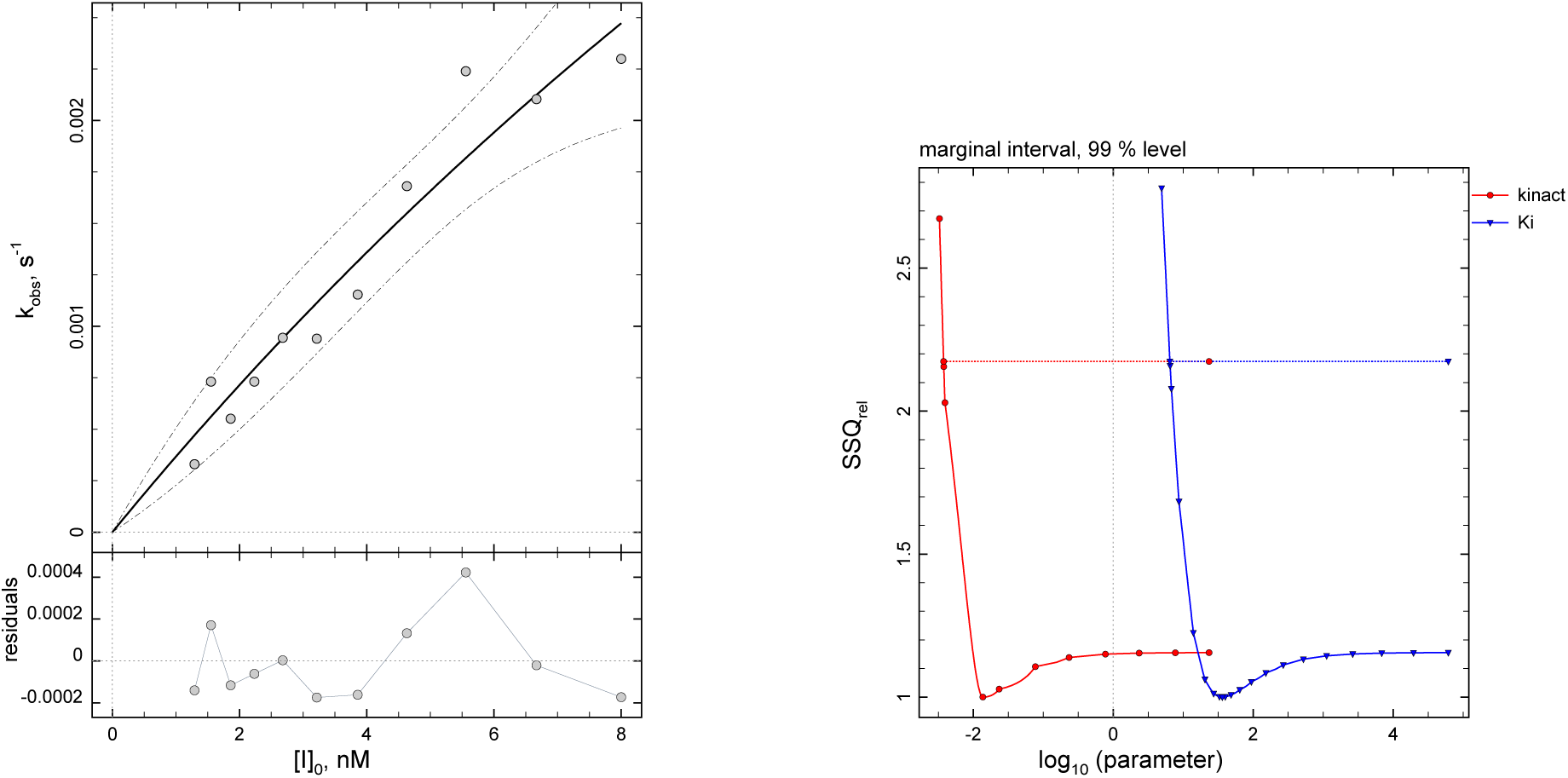
Results of fit to the two-step hyperbolic Eqn (2), data set no. 6. **Left:** Experimental data (circles), best-fit model curve, and the residual plot. **Right:** Confidence interval profiles for the kinetic constants 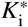 and *k*_inact_.

### 3.2. Akaike and Bayesian information criteria

The results of statistical model discrimination based on the Akaike Information Criterion are summarized in *Table 4*. The column labeled *S* _1_/*S* _2_ represents the ratio of sum of squares obtained for the one-step linear fitting Eqn (3) (*S* _1_) divided by the sum of squares obtained for the two-step hyperbolic fitting Eqn (3) (*S* _2_). As is expected from theory, the linear fitting equation always produced a larger sum of squares than the hyperbolic fitting equation, simply because the hyperbolic equation Eqn (3) contains an additional adjustable parameter.

**Table 4:** Results of model discrimination analysis based on the Akaike information criterion AIC. For details see text.

| set no. | $S_1/S_2$ | $\Delta AIC$ | $w_{AIC}$ | model<sub>95%</sub> | model<sub>99%</sub> |
| --- | --- | --- | --- | --- | --- |
| 1 | 1.7 | 0 | 0.70 | C | C |
| 2 | 1.7 | 0 | 0.69 | C | C |
| 3 | 10.9 | 0 | 1.00 | B | B |
| 4 | 12.7 | 0 | 1.00 | B | B |
| 5 | 1.1 | 3.4 | 0.15 | C | C |
| 6 | 1.2 | 2.3 | 0.24 | C | C |
| 7 | 1.2 | 1.9 | 0.28 | C | C |
| 8 | 2.6 | 0 | 0.96 | B | C |

The column labeled ∆AIC in *Table 4* represents the differential AIC value for the two-step fitting model represented by Eqn (2), corresponding to the two step mechanism **B**. The column labeled *w*_AIC_ represents the corresponding Akaike weight. The columns labeled model_95%_ and model_99%_ in *Table 4* contain the results of a decision in favor of either the two-step inhibition mechanism **B** or, alternately, the one-step inhibition mechanism **C**. The decision was based on whether or not the value in the Akaike weight for the two-step model is grater than 0.95 or 0.99, respectively.

The results show that the two-step inhibition mechanism **B** is favored consistently (across both replicated experiments and at both probability levels) only for acalabrutinib inhibition of BTK. In the case of data set no. 8 (acalabrutinib inhibition of TEC, second replicate), the two-step mechanism was favored at 95% probability level but not at the more stringent 99% probability level. In every other instance the one-step inhibition mechanism **C** was the preferred theoretical model. The best-fit values of the Bayesian Information Criterion, BIC, produced to exactly identical conclusions as those described above for the Akaike Information Criterion.

### 3.3. F-test for nested models

The results based on the *F*-test for nested models are summarized in *Table 5*. Larger values of *F*_test_ signify stronger support to the two-step inhibition mechanism **B**. The ∆*F* values represent the difference *F*_test_ − *F*_crit_ at the given probability level. The corresponding critical values were *F*_0.01_(1, 9) = 10.04 and *F*_0.05_(1, 9) = 4.96. At each probability level (95% or 99%), the two-step mechanism **B** is preferred when ∆*F* > 0, meaning that the *F*_test_ value exceeds the critical value *F*_crit_.

**Table 5:** Results of model discrimination analysis based on *F*-test for nested models. For details see text.

| set no. | $F_{test}$ | $\Delta F_{99\%}$ | model<sub>99%</sub> | $\Delta F_{95\%}$ | model<sub>95%</sub> |
| --- | --- | --- | --- | --- | --- |
| 1 | 6.7 | -3.4 | C | 1.7 | B |
| 2 | 6.6 | -3.5 | C | 1.6 | B |
| 3 | 99.4 | 89.4 | B | 94.5 | B |
| 4 | 117.1 | 107.1 | B | 112.2 | B |
| 5 | 0.5 | -9.6 | C | -4.5 | C |
| 6 | 1.6 | -8.5 | C | -3.4 | C |
| 7 | 2.1 | -8.0 | C | -2.9 | C |
| 8 | 15.7 | 5.6 | B | 10.7 | B |

The results show that unambiguous conclusions could be reached for only two out of four enzyme–inhibitor combinations. First, in the case of acalabrutinib inhibition of BTK, the two-step mechanism **B** is strongly preferred for both replicated data sets and at both probability levels (95% and 99%). Second, in the case of ibrutinib inhibition of TEC, the one-step mechanism **C** is strongly preferred for both replicated data sets and at both probability levels.

In contrast, ibrutinib inhibition of BTK resulted in ambiguous results, because at 99% probability level the one-step mechanism **C** is favored, whereas at 95% probability level the two-step mechanism **B** is favored, however weakly. Another type of ambiguity was encountered for acalabrutinib inhibition of TEC. In that case, on the basis of the first replicate (data set no. 7) the one-step mechanism **C** is quite strongly favored, whereas on the basis of the second replicate (data set no. 8) the opposite is true.

## 4. Discussion

Ever since the discovery of acetylsalicylate (Aspirin) in the 19th century, irreversible enzyme inhibitors continue to be highly important as potential therapeutic agents [16, 17]. Abdeldayem *et al.* [18] recently reviewed ten years of progress specifically in the field of protein kinases as targets, where irreversible inhibitors continue to emerge as viable therapeutics. However, rigorous evaluation of inhibitory potency of irreversible drug candidates presents a number of technical and conceptual challenges, because the overall potency of irreversible inhibitors consists of two separate components. First, the compound’s initial binding affinity (measured by the inhibition constant *K*_i_) is responsible for the reversible formation of a noncovalent complex. Second, the inhibitor’s chemical reactivity (measured by the inactivation rate constant *k*_inact_) determines how rapidly the initial complex is converted to the final covalent conjugate.

It has been proposed that the one-step inhibition mechanism **C** is characteristic for irreversible enzyme inhibitors that have extremely low initial binding affinity and a complete lack of specificity [2, sec. 7.2.1] [3, sec 9.1]. Here we report on the basis of previously published experimental data [4] that ibrutinib, a highly specific inhibitor presumably honing in on the active site of certain protein kinases, inhibits the TEC enzyme apparently in a single step. If a reversibly formed noncovalent complex is actually present, apparently its formation is undetectable by the *k*_obs_ method, in the sense that the plot of *k*_obs_ vs. the inhibitor concentration [I]_0_ is essentially linear (see *Figure 3*).

A commonly encountered reason for the plot of *k*_obs_ vs. [I]_0_ being apparently linear is the fact that the maximum inhibitor concentration used in the assay might not be high enough to be sufficiently informative about the 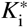 value. Eqn (2) describes a rectangular hyperbola. However, if all inhibitor concentrations utilized in any given assay happen to be significantly lower than the half-saturating point, which is by definition identical to the 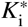 value, the hyperbolic Eqn (2) essentially turns into a linear equation with slope equal to the ratio 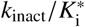. This was pointed out already by Kitz & Wilson [8] in the original paper describing *k*_obs_ method: “If 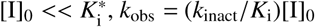 and *k*_inact_/*K*_i_ can be set equal to *k*_eff_; i.e. the kinetics are then not distinguishable from a simple bimolecular mechanism.” Geometrically speaking, it is as though all experimental [I]_0_ values were located in the “initial linear portion” of the hyperbolic curve.

The key observation is that it is not feasible to increase the inhibitor concentration arbitrarily, because the rate of covalent inactivation inevitably increases with the concentration of the inhibitor. At a certain sufficiently high inhibitor concentration, which might still be significantly lower than the 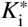, the inactivation reaction might become so fast that the enzyme is nearly fully inhibited before the first time-point is even recorded. In other words, the chemical reactivity of the initial noncovalent complex might be so high that the complex does not exist long enough for us to reliably determine the inhibitor’s noncovalent binding affinity. More precisely, if the inactivation rate constant *k*_2_ ≡ *k*_inact_ happens to be higher than the dissociation rate constant *k*_−1_ in *Figure 1*, the noncovalent complex will be pulled through the overall reaction path so rapidly that it will not make a detectable appearance in the reaction mixture. Under those circumstances, the experimental data might appear as though there were only a single irreversible step (mechanism **C**).

There also exists a fundamental theoretical reason why the *k*_obs_ method might provide grossly misleading results. That reason is rooted in the simplifying assumptions that were used to derive Eqn (1). The mathematical model for the reaction progress of a covalent enzyme assay was derived by Kitz & Wilson [8] under the standard *rapid equilibrium approximation* in enzyme kinetics [19]. Kitz & Wilson assumed that the microscopic rate constant for the inactivation step (i.e., *k*_2_ in mechanism **A**, see *Figure 1*) is negligibly small when compared with the dissociation rate constant of the noncovalent complex (*k*_−1_ in mechanism **A**). As was pointed out by Cornish-Bowden [2, sec 7.2.2], “if *k*_2_ is not small enough to allow formation of E − I to be treated as an equilibrium, […] the loss of activity does not follow simple first-order kinetics; there is no exact analytical solution, but the kinetics may still be analyzed by numerical methods.” Thus, if the inactivation rate constant *k*_2_ happens to be comparable in magnitude with the dissociation rate constant *k*_−1_, the time course of the enzyme reaction does not actually follow Eqn (1) and therefore all *k*_obs_ values determined from it are by definition invalid.

Of course, without performing highly advanced rapid-kinetic measurements we cannot possibly know in advance what is the relationship between *k*_2_ and *k*_−1_ in any given case. However, it is possible to make educated guesses. Let us assume for the sake of discussion that the results obtained on the basis of the *k*_obs_ method for the ibrutinib vs. TEC system are actually valid. The reported values [4, Tab. 1] are *k*_inact_ = 0.013 s^−1^; *K*_i_ = 1.8 × 10^−9^ M and *k*_inact_/*K*_i_ = 8 × 10^6^ M^−1^s^−1^. The inactivation rate constant *k*_inact_ is by definition equal to *k*_2_ in *Figure 1*. According to the steady-state approximation introduced by Malcolm & Radda [20], the covalent inhibition constant is defined as *K*_I_ = (*k*_−1_ + *k*_2_)/*k*_1_. Finally, let us take into account that the catalytic efficiency constant *k*_eff_ ≡ *k*_inact_/*K*_I_ represents the lower limit estimate for the second-order association rate constant *k*_1_ [2]. Under these assumptions, we can obtain an approximate estimate of the dissociation rate constant *k*_−1_, starting from the steady-state definition of *K*_I_, as *k*_−1_ = *K*_I_ ×*k*_1_ −*k*_2_. In this case, *k*_−1_ = 1.8 × 10^−9^ × 8 × 10^6^ − 0.013 = 0.0014 s^−1^. Thus, based on the results reported for the ibrutinib vs. TEC system, the lower limit estimate of the dissociation rate constant *k*_−1_ = 0.0014 s^−1^ is approximately ten times lower than the observed inactivation rate constant *k*_2_ = 0.013 s^−1^. But recall that according to the rapid-equilibrium approximation [19], on which the standard *k*_obs_ method is theoretically based, the rate constant *k*_2_ (inactivation) is supposed to be negligibly small compared to *k*_−1_ (dissociation). Clearly this assumption is violated. Therefore, in the specific case of ibrutinib inhibition of the TEC kinase enzyme, Eqn (1) is almost certainly not a suitable fitting model specifically under the *rapid-equilibrium* approximation.

One possible remedy is to recast Eqn (2) such that the dissociation equilibrium constant *K*_i_ is replaced by the steady-state inhibition constant *K*_I_, although this formal solution introduces its own conceptual challenges discussed in detail by Cornish-Bowden [21]. Another possible solution is to follow Cornish-Bowden’s advice, which essentially suggests abandoning all simplistic algebraic models for the analysis of irreversible inhibition data and rely instead on the fully general “numerical methods” [2, sec 7.2.2]. These numerical methods are based on iteratively solving systems of simultaneous first-order ordinary differential equations (ODEs) and, as such, make no simplifying assumptions about the relative magnitude of microscopic rate constants that appear in any given inhibition mechanisms. The software package DynaFit [5, 6] implements the highly advanced numerical algorithm LSODE (Livermore Solver of Ordinary Differential Equations) [22, 23] and has been used profitably in the study of irreversible inhibition kinetics [24] without any simplifying assumptions.

## 5. Summary and Conclusions

Previously published [4] experimental *k*_obs_ values, generated by fitting reaction progress curves from covalent inhibition assays to Eqn (1), were re-analyzed using four independent statistical criteria. Contrary to the conclusion of the original report [4], the two-step mechanism **B** could be firmly established only for acalabrutinib inhibition of BTK, as one of the four possible enzyme/inhibitor pairs. In this case, the best-fit values of *K*_i_ and *k*_inact_ are fully supported by the data. In the three remaining enzyme/inhibitor combinations the mechanistic conclusions differ from the originally published results. In the specific case of ibrutinib inhibition of TEC, there is virtually no support for the two-step mechanism, whereas in the cases of ibrutinib inhibition of BTK and acalabrutinib of TEC the results are ambiguous. In these three cases, only the ratio *k*_inact_/*K*_i_ and the lower limit estimates for the individual values of *K*_i_ and *k*_inact_ can be known with high degree of certainty.

The model-selection methods, newly presented in this report as a unified statistical toolkit, should prove useful in other research projects that also involve the mechanistic analysis of *k*_obs_ values arising in the study of covalent inhibition kinetics. The software package DynaFit [6], utilized for all data analyses contained in this report, is available free of charge to all academic institutions as a download from http://www.biokin.com.

## Acknowledgements

I thank Dr. Claire McWhirter (Artios Pharma, Cambridge, UK) for helpful discussions.

## Supporting information

1. Raw data files (*k*_obs_ vs. [I]_0_) for data sets no. 1–6 listed in *Table 1*.
2. Listing of DynaFit [6] script files that were used to generate this report.
3. Detailed step-by-step instructions on how to use the DynaFit software package to repeat the data analyses reported here.

## Appendix

### A. Explanation of symbols

**Table A.1:** Explanation of symbols: Microscopic rate constants and derived kinetic constants.

| Symbol | Unit | Explanation |
| --- | --- | --- |
| $k_{1s}$ | $M^{-1}s^{-1}$ | association rate constant for $E+S \rightarrow E \cdot S$ |
| $k_{-1s}$ | $s^{-1}$ | dissociation rate constant for $E \cdot S \rightarrow E + S$ |
| $k_{2s}$ | $s^{-1}$ | turnover number; $k_{2s} \equiv k_{cat}$ |
| $K_M$ | M | Michaelis constant; $K_M = (k_{-1s} + k_{2s})/k_1$ |
| $k_S$ | $M^{-1}s^{-1}$ | catalytic efficiency constant; specificity constant; $k_S \equiv k_{cat}/K_M$ |
| $k_1$ | $M^{-1}s^{-1}$ | association rate constant for $E+I \rightarrow \dots$ |
| $k_{-1}$ | $s^{-1}$ | dissociation rate constant for $E \cdot I \rightarrow E + I$ |
| $k_2$ | $s^{-1}$ | inactivation rate constant; $k_2 \equiv k_{inact}$ |
| $k_1^*$ | $M^{-1}s^{-1}$ | apparent association rate constant:<br>competitive: $k_1^* = k_1/(1 + [S]_0/K_M)$<br>uncompetitive: $k_1^* = k_1 (1 + [S]_0/K_M)$<br>noncompetitive $k_1^* = k_1$ |
| $k_{eff}$ | $M^{-1}s^{-1}$ | second-order inhibition efficiency constant; $k_{eff} \equiv "k_{inact}/K_i"$:<br>steady-state two-step mechanism **A**: $k_{eff} = k_1 k_2/(k_{-1} + k_2)$<br>rapid-equilibrium two-step mechanism **B**: $k_{eff} = k_1 k_2/k_{-1}$<br>one-step mechanism **C**: $k_{eff} = k_1$ |
| $k_{eff}^*$ | $M^{-1}s^{-1}$ | apparent inhibition efficiency constant:<br>competitive: $k_{eff}^* = k_{eff}/(1 + [S]_0/K_M)$<br>uncompetitive: $k_{eff}^* = k_{eff} (1 + [S]_0/K_M)$<br>noncompetitive $k_{eff}^* = k_{eff}$ |
| $K_i$ | M | equilibrium dissociation constant of the $E \cdot I$ complex; $K_i = k_{-1}/k_1$ |
| $K_i^*$ | M | apparent equilibrium dissociation constant:<br>competitive: $K_i^* = K_i (1 + [S]_0/K_M)$<br>uncompetitive: $K_i^* = K_i/(1 + [S]_0/K_M)$<br>noncompetitive $K_i^* = K_i$ |
| $K_I$ | M | inhibition constant; $K_I = (k_{-1} + k_2)/k_1$ |
| $K_I^*$ | M | apparent inhibition constant:<br>competitive: $K_I^* = K_I (1 + [S]_0/K_M)$<br>uncompetitive: $K_I^* = K_I/(1 + [S]_0/K_M)$<br>noncompetitive $K_I^* = K_I$ |
| $k_{obs}$ | $s^{-1}$ | apparent first-order rate constant for enzyme inactivation |

**Table A.2:** Explanation of symbols: Concentrations, reaction rates, and auxiliary symbols.

| Symbol | Unit | Explanation |
| --- | --- | --- |
| $[X]$ | M | concentration of reactant X, where X = S, P, I, or E |
| $[X]_0$ | M | initial (total, analytic) concentration of reactant X |
| $F$ | AIU | observed experimental signal in arbitrary instrument units (AIU) |
| $F_0$ | AIU | baseline signal; baseline offset |
| $r_P$ | AIU/M | molar response coefficient of the reaction product P |
| $v_0$ | $\text{Ms}^{-1}$ | initial rate of the uninhibited enzyme reaction, at $[I]_0 = 0$ |
| $V_0$ | $\text{AIU s}^{-1}$ | observed uninhibited initial rate in arbitrary instrument units |
| $v_i$ | $\text{Ms}^{-1}$ | initial rate of the inhibited enzyme reaction, at $[I]_0 > 0$ |
| $V_i$ | $\text{AIU s}^{-1}$ | observed inhibited initial rate in arbitrary instrument units |
| $\alpha, \beta, \gamma$ | $\text{s}^{-1}$ | auxiliary variables (groupings of rate constants) |
| $r_1, r_2$ | $\text{s}^{-1}$ | apparent bi-exponential rate constants |
| $a_1, a_2$ | – | bi-exponential amplitudes |

## References

[1] S. Krishnan, R. M. Miller, B. Tian, R. D. Mullins, M. P. Jacobson, J. Taunton, Design of reversible cysteine-targeted michael acceptors guided by kinetic and computational analysis 136 (2014) 12624–12630. URL https://doi.org/10.1021/ja505194w

[2] A. Cornish-Bowden, Fundamentals of Enzyme Kinetics, 4th Edition, Wiley-VCH Verlag, Berlin, 2012.

[3] R. A. Copeland, Evaluation of Enzyme Inhibitors in Drug Discovery, 2nd Edition, John Wiley, New York, 2013.

[4] M. Hopper, T. Gururaja, T. Kinoshita, J. P. Dean, R. J. Hill, A. Mongan, Relative selectivity of irreversible inhibitors requires assessment of inactivation kinetics and cellular occupancy: A case study of ibrutinib and acalabrutinib, J. Pharm. Exp. Therap. 372 (2020) 331–338. URL https://doi.org/10.1124/jpet.119.262063

[5] P. Kuzmič, Program DYN AFIT for the analysis of enzyme kinetic data: Application to HIV proteinase, Anal. Biochem. 237 (1996) 260–273. URL http://doi.org/10.1006/abio.1996.0238

[6] P. Kuzmič, DynaFit - A software package for enzymology, Meth. Enzymol. 467 (2009) 247–280. URL http://doi.org/10.1016/S0076-6879(09)67010-5

[7] P. Kuzmič, A steady-state algebraic model for the time course of covalent enzyme inhibition, BioRxiv (2020) doi:10.1101/2020.06.10.144220. URL https://doi.org/10.1101/2020.06.10.144220

[8] R. Kitz, I. B. Wilson, Esters of methanesulfonic acid as irreversible inhibitors of acetyl-cholinesterase, J. Biol. Chem. 237 (1962) 3245–3249. URL https://www.jbc.org/content/237/10/3245.long

[9] W. X. Tian, C. L. Tsou, Determination of the rate constant of enzyme modification by measuring the substrate reaction in the presence of the modifier, Biochemistry 21 (1982) 1028–1032. URL https://doi.org/10.1021/bi00534a031

[10] D. M. Bates, D. G. Watts, Nonlinear Regression Analysis and its Applications, Wiley, New York, 1988.

[11] D. G. Watts, Parameter estimation from nonlinear models, Meth. Enzymol. 240 (1994) 24–36. URL https://doi.org/10.1016/S0076-6879(94)40041-5

[12] K. B. Burnham, D. R. Anderson, Model Selection and Multimodel Inference: A Practical Information-Theoretic Approach, 2nd Edition, Springer-Verlag, New York, 2002.

[13] J. I. Myung, M. A. Pitt, Model comparison methods, Meth. Enzymol. 383 (2004) 351–366. URL https://doi.org/10.1016/S0076-6879(04)83014-3

[14] J. I. Myung, Y. Tang, M. A. Pitt, Evaluation and comparison of computational models, Meth. Enzymol. 454 (2009) 287–304. URL https://doi.org/10.1016/S0076-6879(08)03811-1

[15] B. Mannervik, Regression analysis, experimental error, and statistical criteria in the design and analysis of experiments for discrimination between rival kinetic models, Meth. Enzymol. 87 (1982) 370–390. URL https://doi.org/10.1016/S0076-6879(82)87023-7

[16] Z. Zhao, P. E. Bourne, Progress with covalent small-molecule kinase inhibitors, Drug Disc. Today 23 (2018) 727–735. URL https://doi.org/10.1016/j.drudis.2018.01.035

[17] A. K. Ghosh, I. Samanta, A. Mondal, W. R. Liu, Covalent inhibition in drug discovery, ChemMedChem 14 (2019) 889–906. URL https://doi.org/10.1002/cmdc.201900107

[18] A. Abdeldayem, Y. Raouf, S. Constantinescu, R. Moriggl, P. Gunning, Advances in covalent kinase inhibitors, Chem. Soc. Rev. 49 (2020) 2617–2687. URL https://doi.org/10.1039/c9cs00720b

[19] I. H. Segel, Enzyme Kinetics: Behavior and Analysis of Rapid Equilibrium and Steady-state Enzyme Systems, Wiley, New York, 1975.

[20] A. D. B. Malcolm, G. K. Radda, The reaction of glutamate dehydrogenase with 4-iodoacetarnido salicylic acid, Eur. J. Biochem. 15 (1970) 555–561. URL http://doi.org/10.1111/j.1432-1033.1970.tb01040.x

[21] A. Cornish-Bowden, Validity of a “steady-state” treatment of inactivation kinetics, Eur. J. Biochem. 93 (1979) 383–385. URL https://doi.org/10.1111/j.1432-1033.1979.tb12834.x

[22] A. C. Hindmarsh, LSODE and LSODI, two new initial value ordinary differential equation solvers, ACM SIGNUM Newslett. 15 (1980) 10–11. URL https://doi.org/10.1145/1218052.1218054

[23] A. C. Hindmarsh, ODEPACK: a systematized collection of ODE solvers, in: R. S. Stepleman, M. Carver, R. Peskin, W. F. Ames, R. Vichnevetsky (Eds.), Scientific Computing, North Holland, Amsterdam, 1983, pp. 55–64.

[24] P. A. Schwartz, P. Kuzmicč, J. Solowiej, S. Bergqvist, B. Bolanos, C. Almaden, A. Nagata, K. Ryan, J. Feng, D. Dalvie, J. C. Kath, M. Xu, R. Wani, B. W. Murray, Covalent EGFR inhibitor analysis reveals importance of reversible interactions to potency and mechanisms of drug resistance, Proc. Natl. Acad. Sci. U.S.A. 111 (2014) 173–178. URL https://doi.org/10.1073/pnas.1313733111

