## Supporting Information for "Deciding between one-step and two-step irreversible inhibition mechanisms on the basis of “*k*_obs_” data: A statistical approach"

Petr Kuzmič

*BioKin Ltd., Watertown, Massachusetts, USA*
*http://www.biokin.com*

---

---

**References** **8**

### 1. DynaFit script file listing

The DynaFit [1] script that was used to analyze the experimental $k_{obs}$ data listed in Supplemental Table S1 in ref. [2] is listed below. Please refer to the *DynaFit Scripting Manual* available from www.biokin.com for a detailed explanation of the syntax and semantics.

```
Hopper et al. (2020) J. Pharmacol. Experim. Ther. 372, 331-338.
Fit kobs vs. [I] data from Supplemental Table 1
;_______________________________________________________________________
[task]
   task = fit
   data = generic
   model = two-step ?
[parameters]
   Io, kinact, Ki
[model]
   kinact = 0.001 ??
   Ki = 10 ??
   kobs = kinact * Io / (Io + Ki)
[data]
   variable Io
   directory ./technotes/2020/01/data
   sheet T1.csv
   column 2
[output]
   directory ./technotes/2020/01/output/T1-99
[settings]
{Output}
   XAxisLabel = [I]_0, nM
   YAxisLabel = k_{obs}, s^{-1}
   ConfidenceBands = y
   PredictionBands = n
{ConfidenceIntervals}
   LevelPercent = 99
{ModelSelection}
   NestedModels = y
   FCriticalLevelPercent = 99
;_______________________________________________________________________
[task]
   task = fit
   data = generic
   model = one-step ?
[parameters]
   Io, keff
[model]
   keff = 0.000001 ??
   kobs = keff * Io
[end]
;_______________________________________________________________________
;_______________________________________________________________________
```

### 2. DynaFit instructions

This section provides step-by-step instructions on how to repeat the data analyses describe in the main manuscript by using the software package DynaFit.

#### *2.1. Downloading and installing DynaFit*

To download and install the DynaFit software package [1], please follow these steps:

1. Point a web browser to www.biokin.com/dynafit/download.html
2. Click the [Download] link
3. Save the downloaded file dynafit4-win.exe anywhere in the MS Windows user space (for example, "Documents").
4. Use MS File Explorer to navigate to the destination directory, e.g. Documents.
5. Double-click on file dynafit4-win.exe.
6. In the "Open File – Security Warning" dialog, click [Run]
7. Select the final destination directory, e.g. C:/Users/USERNAME/Documents.
8. Navigate into the newly created subdirectory DynaFit4.
9. Double-click on the MS Widows Command File SetDynaFifPath.bat. The message displayed in the command window should be similar to this:

```
SUCCESS: Specified value was saved.
DynaFit path was set to [C:/Users/USERNAME/Documents/DynaFit4]
```

10. Right-click on the file DynaFit.exe in the subdirectory DynaFit4.
11. Select menu item *Send to ... Desktop (Create Shortcut).*

Note that the DynaFit software package does *not* use the standard MS Windows installer. In this way the software can be installed successfully even in highly restrictive corporate or academic environments, for example, when strict IT administration policies do not allow end-users to install software packages in the standard way. After this installation procedure is completed, the GUI (Graphical User Interface) version of the DynaFit software package can be triggered by using the newly created desktop shortcut. To verify that the installation was successful, please follow the steps:

1. Start DynaFit.
2. Select menu item *View ... Startup Path.* The message displayed in the output window should be similar to this:

```
DynaFit startup path:

C:\Users\Petr\Documents\DynaFit4
```

The displayed DynaFit startup path must *not* contain C:/Windows, if it does, it means that DynaFit was started by using the Windows 10 "Search App" system. DynaFit will not operate properly when started by Windows 10 "Search App". It must be started by double-clicking on the file DynaFit.exe or on the shortcut to it.

#### 2.2. *Analyzing the $k_{obs}$ data*

Assuming that the DynaFit installation was successful (see section 2.1), the $k_{obs}$ vs. $[I]_0$ data initially published in ref. [2] can be analyzed as follows:

1. Start MS File Explorer.
2. Navigate to the following subdirectory:
   DynaFit4/DynaFit/technotes/2020/01
3. Double-click on the following file:
   DynaFit4/DynaFit/technotes/2020/01/RunTechNote.bat
4. Press the [Enter] key
5. Navigate in the output files that should automatically open in the default web browser.

### 3. Raw data

#### 3.1. *Dataset numbering scheme*

Dataset numbering utilized in this report is displayed in *Table S1*. BTK is Bruton tyrosine kinase; TEC is tyrosine kinase expressed in hepatocellular carcinoma. The numbering of datasets (1 – 8) corresponds to Supplemental Tables 1 – 8 and Supplemental Figures 1 – 8 originally published in [2].

| dataset no. | enzyme | $[E]_0$ | inhibitor |
|---|---|---|---|
| 1 | BTK | 0.35 | ibrutinib |
| 2 | BTK | 0.045 | ibrutinib |
| 3 | BTK | 0.35 | acalabrutinib |
| 4 | BTK | 0.045 | acalabrutinib |
| 5 | TEC | 0.3 | ibrutinib |
| 6 | TEC | 0.115 | ibrutinib |
| 7 | TEC | 0.3 | acalabrutinib |
| 8 | TEC | 0.2 | acalabrutinib |

**Table S1:** Dataset numbering utilized in this report.

#### 3.2. *Raw experimental data*

##### 3.2.1. *Data set no. 1*

| $[I]_0$ nM | $k_{obs}$ $s^{-1}$ | error |
|---:|---:|---:|
| 10 | 0.002622204 | 0.000360793 |
| 8.333333333 | 0.002546468 | 0.000330995 |
| 6.944444444 | 0.002405458 | 0.000221326 |
| 5.787037037 | 0.001744919 | 0.000127889 |
| 4.822530864 | 0.00177236 | 0.000185465 |
| 4.01877572 | 0.001146117 | 0.000100137 |
| 3.348979767 | 0.000984421 | 4.18082E-05 |
| 2.790816472 | 0.000909377 | 3.23242E-05 |
| 2.325680394 | 0.000821419 | 2.85424E-05 |
| 1.938066995 | 0.00067625 | 2.23957E-05 |
| 1.615055829 | 0.000610541 | 4.48376E-05 |

**Table S2:** Data set no. 1 from Supplemental Table 1 of ref. [2].

##### 3.2.2. *Data set no. 2*

| $[I]_0$ nM | $k_{obs}$ $s^{-1}$ | error |
|---:|---:|---:|
| 5 | 0.001745425 | 0.000253643 |
| 4.166666667 | 0.001578286 | 0.000140563 |
| 3.472222222 | 0.00126266 | 0.000073267 |
| 2.893518519 | 0.001018874 | 6.78472E-05 |
| 2.411265432 | 0.000931715 | 4.25298E-05 |
| 2.00938786 | 0.000808285 | 3.47423E-05 |
| 1.674489883 | 0.000656164 | 2.93531E-05 |
| 1.395408236 | 0.00054087 | 2.39779E-05 |
| 1.162840197 | 0.000468612 | 1.93592E-05 |
| 0.969033497 | 0.000317853 | 1.73319E-05 |
| 0.807527914 | 0.000365215 | 1.46379E-05 |

**Table S3:** Data set no. 2 from Supplemental Table 2 of ref. [2].

In the original publication [2], the $[I]_0$ = 0.969033497 data point was deleted. In this report, all data points were analyzed.

*3.2.3. Data set no. 3*

| $[I]_0$ nM | $k_{obs}$ $s^{-1}$ | error |
|---:|---:|---:|
| 321.97 | 0.00097791 | 3.61277E-05 |
| 268.3083333 | 0.000956686 | 3.22938E-05 |
| 223.5902778 | 0.000907469 | 3.34095E-05 |
| 186.3252315 | 0.000846832 | 2.59687E-05 |
| 155.2710262 | 0.000714928 | 2.19069E-05 |
| 129.3925219 | 0.000692553 | 3.28869E-05 |
| 107.8271016 | 0.000487165 | 0.000019903 |
| 89.85591796 | 0.000462368 | 1.87039E-05 |
| 74.87993163 | 0.000420859 | 1.36238E-05 |
| 62.39994303 | 0.000367365 | 1.18944E-05 |
| 51.99995252 | 0.000290036 | 7.58E-06 |

**Table S4:** Data set no. 3 from Supplemental Table 3 of ref. [2].

*3.2.4. Data set no. 4*

| $[I]_0$ nM | $k_{obs}$ $s^{-1}$ | error |
|---:|---:|---:|
| 178.9 | 0.000997718 | 5.57243E-05 |
| 149.0833333 | 0.000849673 | 5.84386E-05 |
| 124.2361111 | 0.000775032 | 4.21489E-05 |
| 103.5300926 | 0.000655515 | 3.59637E-05 |
| 86.27507716 | 0.000631197 | 3.11489E-05 |
| 71.89589763 | 0.000496656 | 2.36196E-05 |
| 59.91324803 | 0.000462513 | 1.61551E-05 |
| 49.92770669 | 0.000367588 | 1.62148E-05 |
| 41.60642224 | 0.000337013 | 1.39989E-05 |
| 34.67201853 | 0.000267423 | 1.35051E-05 |
| 28.89334878 | 0.000258543 | 1.29368E-05 |

**Table S5:** Data set no. 4 from Supplemental Table 4 of ref. [2].

#### 3.2.5. *Data set no. 5*

| $[I]_0$ nM | $k_{obs}$ $s^{-1}$ | error |
|---:|---:|---:|
| 15 | 0.003711357 | 0.000325344 |
| 12.5 | 0.003203723 | 0.000259978 |
| 10.41666667 | 0.002651187 | 0.00019961 |
| 8.680555556 | 0.002183536 | 0.00018315 |
| 7.233796296 | 0.001448589 | 0.000194026 |
| 6.02816358 | 0.001166187 | 6.24796E-05 |
| 5.02346965 | 0.001551147 | 4.94976E-05 |
| 4.186224709 | 0.001399701 | 5.60526E-05 |
| 3.48852059 | 0.001096532 | 6.49649E-05 |
| 2.907100492 | 0.001014172 | 7.17453E-05 |
| 2.422583743 | 0.000728765 | 7.20165E-05 |

**Table S6:** Data set no. 5 from Supplemental Table 5 of ref. [2].

In the original publication [2], the $[I]_0$ = 7.233796296 and $[I]_0$ = 6.02816358 data points were deleted. In this report, all data points were analyzed.

#### 3.2.6. *Data set no. 6*

| $[I]_0$ nM | $k_{obs}$ $s^{-1}$ | error |
|---:|---:|---:|
| 8 | 0.002299512 | 0.000361184 |
| 6.666666667 | 0.002103881 | 0.000225242 |
| 5.555555556 | 0.002239668 | 0.000315757 |
| 4.62962963 | 0.001681009 | 0.00016689 |
| 3.858024691 | 0.001154146 | 8.28214E-05 |
| 3.215020576 | 0.000939698 | 6.22118E-05 |
| 2.679183813 | 0.000944351 | 7.14332E-05 |
| 2.232653178 | 0.000731845 | 0.000149011 |
| 1.860544315 | 0.000551128 | 0.000254758 |
| 1.550453596 | 0.000731739 | 0.000384208 |
| 1.292044663 | 0.000330385 | 0.000190587 |

**Table S7:** Data set no. 6 from Supplemental Table 6 of ref. [2].

#### 3.2.7. *Data set no. 7*

| $[I]_0$ nM | $k_{obs}$ $s^{-1}$ | error |
|---:|---:|---:|
| 804.9 | 0.00235 | 0.000128222 |
| 670.75 | 0.001969861 | 0.000115078 |
| 558.9583333 | 0.001576885 | 9.46784E-05 |
| 465.7986111 | 0.001410129 | 0.000131697 |
| 388.1655093 | 0.00100748 | 8.94748E-05 |
| 323.4712577 | 0.000836627 | 5.11058E-05 |
| 269.5593814 | 0.000978956 | 5.53943E-05 |
| 224.6328179 | 0.000885238 | 5.12951E-05 |
| 187.1940149 | 0.000716575 | 4.57308E-05 |
| 155.9950124 | 0.000642043 | 0.000061225 |
| 129.9958437 | 0.000441482 | 4.64066E-05 |

**Table S8:** Data set no. 7 from Supplemental Table 7 of ref. [2].

#### 3.2.8. *Data set no. 8*

| $[I]_0$ nM | $k_{obs}$ $s^{-1}$ | error |
|---:|---:|---:|
| 457.57 | 0.001630108 | 0.000129663 |
| 381.3083333 | 0.001398252 | 9.42539E-05 |
| 317.7569444 | 0.001188951 | 8.23696E-05 |
| 264.7974537 | 0.000964911 | 0.00005898 |
| 220.6645448 | 0.000789929 | 5.28222E-05 |
| 183.8871206 | 0.000701655 | 3.70468E-05 |
| 153.2392672 | 0.000613403 | 2.93045E-05 |
| 127.6993893 | 0.000519269 | 2.58391E-05 |
| 106.4161578 | 0.000460735 | 2.78285E-05 |
| 88.68013148 | 0.000371238 | 2.65775E-05 |
| 73.90010956 | 0.00032225 | 2.50307E-05 |

**Table S9:** Data set no. 8 from Supplemental Table 8 of ref. [2].

### References

[1] P. Kuzmič, DynaFit - A software package for enzymology, Meth. Enzymol. 467 (2009) 247–280.
URL http://doi.org/10.1016/S0076-6879(09)67010-5

[2] M. Hopper, T. Gururaja, T. Kinoshita, J. P. Dean, R. J. Hill, A. Mongan, Relative selectivity of irreversible inhibitors requires assessment of inactivation kinetics and cellular occupancy: A case study of ibrutinib and acalabrutinib, J. Pharm. Exp. Therap. 372 (2020) 331–338.
URL https://doi.org/10.1124/jpet.119.262063
